# Dissociating the Effects of Light at Night from Circadian Misalignment in a Neurodevelopmental Disorder Mouse Model Using Ultradian Light-Dark Cycles

**DOI:** 10.1101/2025.07.04.663193

**Authors:** Sophia Anne Marie B. Villanueva, Huei-Bin Wang, Kyle Nguyen-Ngo, Caihan Tony Chen, Gemma Stark, Gene D. Block, Cristina A. Ghiani, Christopher S. Colwell

## Abstract

Individuals with neurodevelopmental disorders (NDDs) often experience sleep disturbances and are frequently exposed to light during nighttime hours. Our previous studies using the *Cntnap2* knockout (KO) mouse model of NDDs demonstrated that nighttime light exposure increases behaviors such as excessive grooming, reduces social interactions, and disrupts daily locomotor rhythms. To further evaluate the effects of nighttime light exposure, we exposed wild-type (WT) and *Cntnap2* KO mice to an ultradian lighting cycle (T7), which alternates 3.5 hours of light and 3.5 hours of darkness. Circadian rhythms in activity, corticosterone levels, and clock gene expression are maintained under T7 lighting despite the presence of light during the usual night phase, whilst animals display increased depressive-like behaviors and reduce performance on the novel object recognition test. Based on these observations, we hypothesized that T7 lighting would mimic the impact of nighttime light exposure seen in standard light-dark cycles with dim light at night (DLaN). However, in this study, adult WT and *Cntnap2* KO mice held under the T7 cycle did not show the increased grooming behavior or reduced social interaction observed in *Cntnap2* KO mice exposed to DLaN. Regarding locomotor activity rhythms, the T7 cycle lengthened the circadian period and weakened the rhythm amplitude but did not abolish rhythmicity in either genotype. Finally, opposite to DLaN, neither the T7 cycle nor constant darkness (DD) elicited an increase in cFos expression in the basolateral amygdala in WT and KO mice. These results demonstrate that the adverse behavioral and neurobiological effects of nighttime light exposure in a model of a neurodevelopmental disorder depend on circadian disruption rather than light exposure alone, highlighting the importance of circadian stability as a protective factor in NDDS.

## Introduction

A significant proportion of individuals with neurodevelopmental disorders (NDD) experience disruptions in their daily sleep-wake cycles. Common complaints include delayed bedtimes and frequent nighttime awakenings (Martínez-Cayuelas et al., 2022; Sadikova et al., 2023; Al Lihabi, 2023; Peters et al., 2024). These nighttime sleep difficulties are associated with challenging daytime behaviors for the impacted individuals (Mazurek and Sohl, 2016; Veatch et al., 2017) but also negatively affect the sleep and overall health of their parents/caregivers (Hodge et al., 2013; Lopez-Wagner et al., 2008). Additionally, disrupted sleep patterns may lead to increased nighttime exposure to light from electronic screens (Mazurek et al., 2016; Dong et al., 2021; Nagata et al., 2024), which even in young people without NDD has been shown to delay sleep onset (Chang et al., 2015; Gronli et al., 2016; Didikoglu et al., 2023; Stefanopoulou et al., 2024; Guindon et al., 2024).

This led us to explore whether nighttime light exposure could disrupt circadian timing and exacerbate symptoms associated with NDD, using the *Cntnap2* knockout (KO) mouse model. Variants of the *Cntnap2* gene have been linked with autism spectrum disorders (ASD), intellectual disabilities, and seizures in both humans (Arking et al., 2008; Bakkaloglu et al., 2008; Nord et al., 2011; O’Roak et al., 2011; Strauss et al., 2006) and mouse models (Peñagarikano et al., 2011; Poliak et al., 1999). The validity of the *Cntnap2* KO model is supported by findings from intervention studies showing that, for instance, the FDA-approved drug risperidone reduces repetitive behaviors (Peñagarikano et al., 2011), administration of oxytocin enhances social behaviors (Choe et al., 2022), and melatonin treatment improves sleep-wake rhythms (Wang et al., 2020) in these mutants. The *Cntnap2* KO model on its own also shows disruptions in circadian locomotor activity (Angelakos et al., 2019; Wang et al., 2020) and EEG-defined sleep patterns (Thomas et al., 2017; Bougeard et al., 2024).

We have previously shown that exposing the *Cntnap2* KO mice to dim light at night (DLaN) exacerbated the already disrupted locomotor activity and altered the neural activity in the suprachiasmatic nucleus (SCN), as well as the rhythms in clock gene expression as measured by PER2::LUC (Wang et al., 2000). Notably, DLaN elicited excessive repetitive grooming behaviors otherwise not observed in the *Cntnap2* KO model (Wang et al., 2020; 2023). An unresolved question is whether the detrimental effects of nighttime light exposure are due to light’s direct impact on behaviors or to the circadian disruption driven by DLaN. One approach to investigate this issue is to place the animals on an ultradian light-dark (LD) cycle that falls outside the circadian clock’s entrainment range. Prior work by Hattar and colleagues employed an ultradian 3.5-hour light/3.5-hour dark (T7) cycle to pinpoint the effects of light alone (LeGates et al., 2012; Duy and Hattar, 2017; Fernandez et al., 2018). They reported that while the T7 cycle lengthens the circadian period, it does not disrupt internal rhythmicity of the SCN or causes arrhythmicity in sleep architecture and body temperature. The T7 cycle, however, did induce mood disruption without increasing anxiety, suggesting that nighttime light alone can affect mood.

In the present study, we examined the impact of ultradian light exposure on wild-type (WT) and *Cntnap2* KO mice housed under T7 conditions (LD 3.5/3.5). We measured social interactions using the three-chamber test, assessed repetitive grooming behavior, and examined the impact of the T7 light cycle on locomotor rhythms via passive infrared (PIR) detection. Finally, the effects of the T7 lighting were investigated on the expression of cFos in the basal lateral amygdala (BLA), a region implicated in repetitive behaviors, since we have previously reported a significantly higher number of cells positive for this neural activity marker in the BLA of both WT and mutant mice after two weeks of DLaN (Wang et al., 2023).

## Materials and methods

### Animals & Experimental groups

All animal procedures were performed in accordance with the UCLA animal care committee’s regulations. A total of 111 adult (3-4 months of age) mice (55 WT and 56 mutants) were used for these experiments with a mixed number of males and females in each experimental group. *Cntnap2^tm2Pele^* mutant mice (Poliak et al., 1999) back-crossed into the C57BL/6J background strain were acquired (B6.129(Cg)- *Cntnap2^tm2Pele^* /J) from The Jackson Laboratory (Bar Harbor, ME; https://www.jax.org/strain/017482). Mice of the WT C57BL/6J and of the *Cntnap2* null mutant (KO) strain were from our breeding colony maintained in an approved facility of the Division of Laboratory Animal Medicine at the University of California, Los Angeles (UCLA). The animals were entrained in a 12h:12h LD cycle for two weeks before being randomly assigned to one of the following groups: (A) continuing in the normal LD cycle, (B) releasing in constant darkness (DD), exposure to (C) DLaN (10 lx illumination during lights off) or (D) the T7 (3.5h light: 3.5h dark) ultradian cycle, (**Fig. 1**) for two additional weeks. We used a two-week exposure to DLaN based on prior data showing that this duration is enough to alter our behavioral measures (Wang et al., 2020). The regular light was set at 250 lx as measured at the floor of the animal holding chambers while the DLaN was measured at 10 lx (see **supplemental Fig. 1** for more information about the lighting).

**Fig 1:**
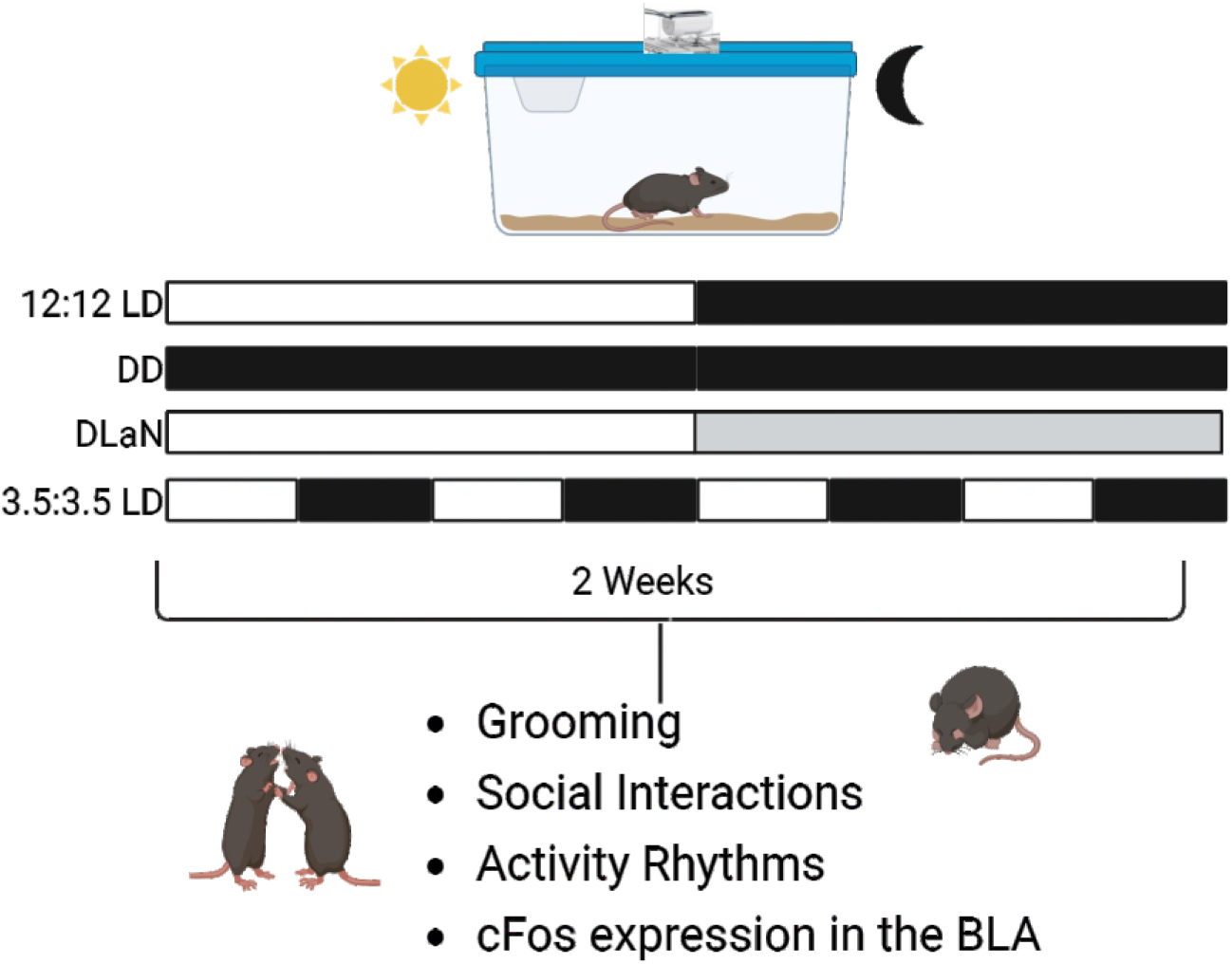
Experimental design. WT and *Cntnap2* KO mice (both sexes) were entrained to a 12h:12h Light-Dark (LD) cycle then randomly assigned to one of four lighting conditions: 12h:12h LD, constant darkness (DD), dim light at night (DLaN) 12 hrs of bright light at 250 lx followed by 12 hrs of dim 10 lx ligh, and a ultradian LD cycle consisting of 3.5 hrs of light followed by 3.5 hrs of dark (T7). Rhythms in cage activity were recorded using passive infrared sensors for the entire time, and at the end of the two weeks, mice were tested at between Zeitgeber time (ZT) or Circadian time (CT) 17-19 for grooming behavior and social interactions. ZT 12 was defined as the time of lights OFF where CT 12 was defined as the time of activity onset when the mice were not synchronized to the external lighting conditions. A separate cohort of mice were exposed to the same 4 lighting conditions and then were euthanized and perfused at ZT18 or CT18. The brains processed for immunofluorescence with an antibody against cFos. Staining in the basolateral Amygdala (BLA) was quantified.

### Behavioral tests

A cohort of 40 WT and 40 *Cntnap2* KO (3-4 months-old) was used to test social and grooming behavior after 2-weeks exposure to one of 4 lighting conditions (LD, DLaN, T7 or DD; **Fig.1**). On the 14^th^ day, the animals were placed in a novel arena for 10 minutes to measure grooming and exploration behavior during their subjective active phase. After habituation to the arena, they were tested using the three-chamber sociability protocol (Yang et al., 2011). In this test, mice are free to roam an arena with three chambers. The central chamber remains empty while the flanking chambers both contain an up-turned metal-grid pencil cup: one is kept empty to be the novel object (O), while an age- and sex-matched WT stranger mouse (S1) is placed in the other. The stranger mice had previously been habituated to the cup for 3 x 15 min sessions. To test for social preference, mice are presented with the choice between O and S1. Social preference is determined by comparing the dwell time of the tested mouse in the two chambers, and the social preference index is calculated using the following formula: (S1 − O)/(S1 + O), where a higher value indicates greater social preference. The time that the tested mouse spent investigating the object or the stranger mouse is also determined. The three-chamber tests were performed under dim red light (<2 lx at arena level) during the night (Zeitgeber time (ZT) 17-19 or Circadian Time (CT) 17-19). Video recordings of each arena were captured using surveillance cameras, supplemented with infra-red lighting, connected to a video-capture card (Adlink Technology Inc., Irvine, CA) on a Dell Optiplex computer system. Mice were automatically tracked using the ANY-maze software (Stoelting, Wood Dale, IL). Grooming and exploration behaviors were manually scored and averaged post-hoc by two independent observers masked to the experimental groups. ANY-maze software determined distance traveled and chamber dwell time during the behavioral tests.

### Cage conditions and activity

Mice were housed individually to monitor and collect locomotor activity rhythms using a top-mounted passive infrared (PIR) motion detector reporting to a VitalView data recording system (Mini Mitter, Bend, OR). The cages were placed in circadian controlled chambers where an environment with a temperature range of 65-75° and humidity levels of 30-40% was maintained. The mice had free access to food and water, and were entrained in a 12h:12h LD cycle for two weeks before being randomly assigned to one of the four experimental groups: LD, DLaN, T7 or DD (**Fig.1**). Each lighting condition was evaluated over two weeks. Detected movements were recorded in 3 min bins, and 14 days of data were averaged for analysis using the Clocklab program (Actimetrics, Wilmette, IL; http://actimetrics.com/products/clocklab/). The strength of the rhythms was determined from the amplitude of the χ2 periodogram at 24 h, to produce the rhythm power (%V) normalized to the number of data points examined. Other locomotor activity parameters were calculated using the Clocklab program. Under LD or DLaN conditions, the time of lights OFF was defined as Zeitgeber time (ZT) 12. Under DD or T7 conditions, the time of activity onset was defined at circadian time (CT) 12.

### ImmunoFluorescence

At the end of the 2-weeks exposure to one of the 4 lighting cycles (**Fig.1**), mice were anesthetized with isoflurane (30%–32%) at a specific time during the night (ZT 18) or subjective night (CT 18) and transcardially perfused with phosphate buffered saline (PBS, 0.1 M, pH 7.4) containing 4% (w/v) paraformaldehyde (Sigma, St. Louis, MO). The brains were rapidly dissected out, post-fixed overnight in 4% PFA at 4 ◦C, and cryoprotected in 15% sucrose. Coronal sections (50 μm) were obtained using a cryostat (Leica, Buffalo Grove, IL), collected sequentially, and paired along the anterior–posterior axis before further processing. Immunohistochemistry was performed as previously described (Lee et al., 2021; Wang et al., 2023). Briefly, free-floating coronal sections containing the BLA were blocked for 1 h at room temperature (1% BSA, 0.3% Triton X-100, 10% normal donkey serum in 1xPBS) and then incubated overnight at 4^◦^C with a rabbit polyclonal antiserum against cFos (1:1000, clone 9F6, Cell Signaling Technology, Danvers, MA) followed by a Cy3-conjugated donkey-anti-rabbit secondary antibody (Jackson ImmunoResearch Laboratories, Bar Harbor, ME). Sections were mounted and coverslips applied with Vectashield mounting medium containing DAPI (4′-6-diamidino-2-phenylinodole; Vector Laboratories, Burlingame, CA), and visualized on a Zeiss AxioImager M2 microscope (Zeiss, Thornwood NY) equipped with a motorized stage, a monochromatic camera AxioCamMRm and the ApoTome imaging system.

### cFos-Positive Cell Counting in the BLA

The BLA was visualized using the DAPI nuclear staining and Z-stack images (7 μm interval, 40 images) of both the left and right BLA were acquired with a 20× objective using the Zeiss Zen digital imaging software. The cells immuno-positive for cFos were counted with the aid of the Zen software tool ‘marker’ in three to five consecutive sections by three observers masked to the genotype and experimental groups. The values obtained from the left and right BLA of each slice were averaged, and the means of the three-five slices were then averaged to obtain one value per animal and are presented as the mean ± S.D. of three to seventeen animals per light treatment.

### Statistical Analysis

Data analysis was performed using Prism (Version 10.5.0; GraphPad Software, La Jolla, CA, USA) or SigmaPlot (version 12.5, SYSTAT Software, San Jose, CA). Two-way Analysis of Variance (ANOVA) followed by the Holm-Sidäk test for multiple comparisons or the Bonferroni’s multiple comparisons test with genotype and light cycle conditions as factors was used to analyse the impact of the different lighting conditions on social interactions and grooming behaviour, parameters of the locomotor activity rhythms and the number of cFos positive cells in the BLA (**Fig. 2**, **3 & 5**; **Tables 1**, **2 & 4**). Three-way ANOVA with sex, genotype, and light cycle conditions as factors was used to analyze the effect of sex on some parameters of locomotor activity rhythms (**Table 3**; **Fig 4**). Values are reported as the mean ± Standard Deviation (SD). Differences were determined to be significant if P <0.05.

**Fig. 2:**
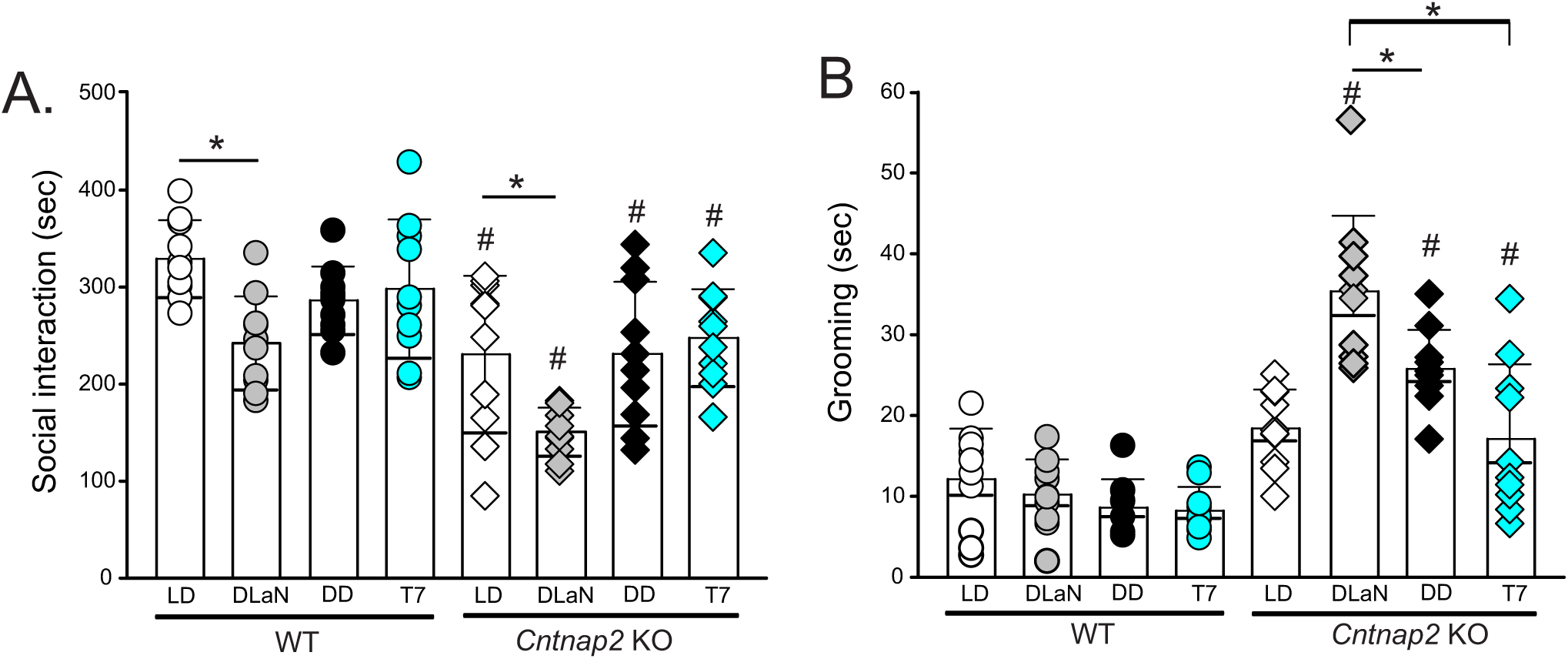
T7 lighting does not mimic DLaN-driven changes in autistic-like behavior in the *Cntnap2* KO mice. A mixed number of male and female mice from both genotypes were held under one of four lighting conditions: LD, DLaN, DD, or a T7 cycle (n= 10 mice per genotype) for two weeks. Behavioral assessments were conducted during the active phase between ZT 17-19 or CT 17-19. (**A**) Social behavior was evaluated by analyzing the time actively spent by the testing mouse interacting with the novel stranger mouse. The stranger and testing mice were matched for age, sex and genotype. DLaN significantly reduced social interactions in both WT and *Cntnap2* KO mice. In contrast, the mice held in T7 did not show changes in social interactions when compared to their counterpart in LD or DD. Time spent interacting with the novel mouse was significantly reduced in the mutants held in any of the light cycle as compared to their respective WT counterpart. (**B**) Grooming was assessed in a novel arena. The time spent grooming was not altered by any of the lighting conditions in WT mice. In contrast, DLaN drove elevated grooming in the *Cntnap2* KO mice. The KO mice held under T7 conditions did not exhibit a significant increase in grooming compared to mutants held in LD or DD conditions. Compared to WT, the *Cntnap2* KO exhibited increased grooming under DLaN, DD, and T7 conditions. Bar graphs show the means ± SD with overlaid the values from individual WT (circles) and *Cntnap2* KO (diamonds) mice held in each of the four lighting conditions: LD (white), DLaN (grey), T7 (teal) and DD (black). Data were analyzed using a two-way ANOVA with genotype and treatment as variables (see **Table 1**) followed by the Holm-Sidak multiple comparisons’ test. Asterisks indicate significant differences (*P < 0.05) between light cycles within genotype, whilst crosshatches indicate significant differences (^#^P < 0.05) between genotypes held in the same lighting condition.

**Fig. 3:**
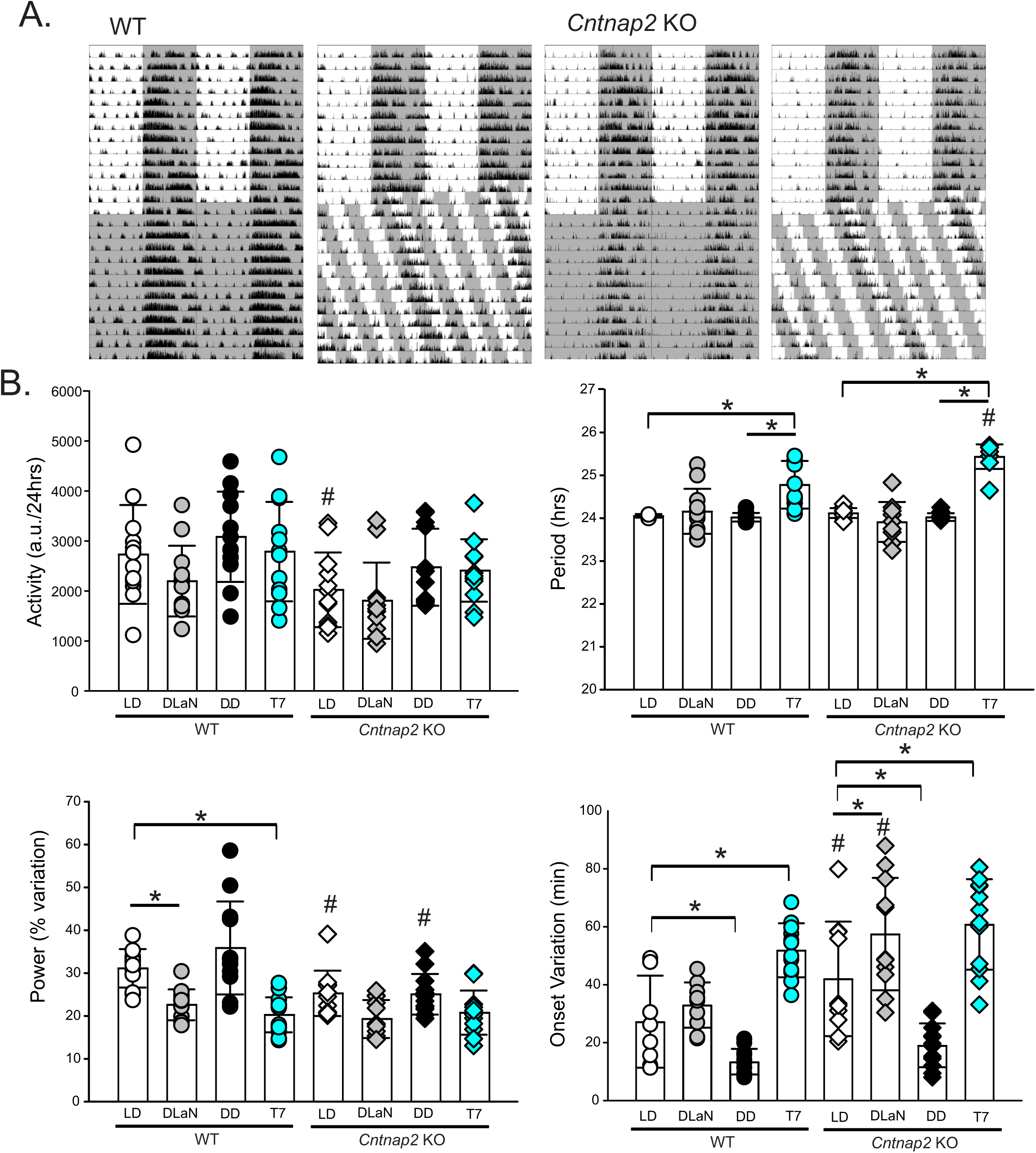
The T7 cycle lengthens the period and weakens, but does not abolish, rhythmicity in locomotor activity. A mixed number of male and female mice of both genotypes were entrained for two weeks in LD and then held for two additional weeks under one of four lighting conditions: LD, DLaN, DD, and a T7 cycle (n= 12 mice per genotype). **(A)** Examples of actograms of daily rhythms in cage activity of WT and *Cntnap2* KO mice held in DD and in a T7 cycle. The activity levels in the actograms were normalized to the same scale (85% of the maximum of the most active individual). Each row represents two consecutive days, and the second day is repeated at the beginning of the next row. The grey shading represents the time of darkness. **(B)** Properties of the daily activity rhythms (10-days recordings) in each of the four lighting conditions: LD (white), DLaN (grey), DD (black), and T7 (teal). In WT mice (circles), the T7 cycle lengthens the period, reduces the power and increases the onset variation of the activity rhythms. In *Cntnap2* KO mice (diamonds), the T7 lighting also increases the period and onset variation. Compared to WT, the KO mice in LD exhibit reduced activity levels, reduced power and increased onset variation (**Table 2**). Bar graphs show the means ± SD with overlaid the values from individual WT (circles) and *Cntnap2* KO (diamonds) mice. Data were analyzed by two-way ANOVA with genotype and treatment as variables (see **Table 2**) followed by the Holm-Sidak multiple comparisons’ test. The asterisks indicate significant differences (*P < 0.05) between lighting cycles within genotype and the crosshatches indicate significant differences (P < 0.05) between the two genotypes under the same lighting cycles.

**Fig. 4:**
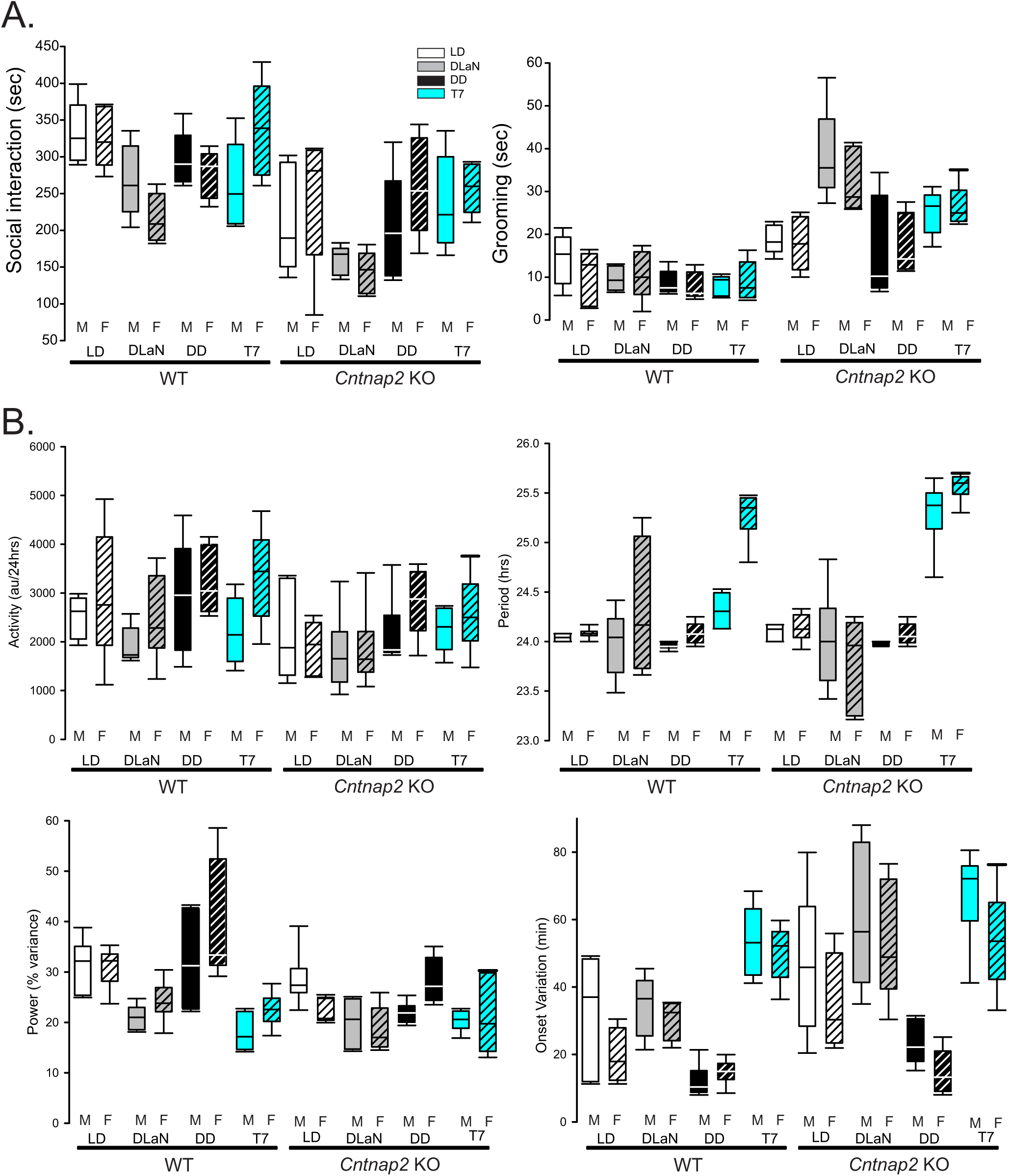
Effect of sex on autistic-like behaviors and activity rhythms in WT and *Cntnap2* KO mice. The behavioral measures described above were also segregated by sex. **(A)** Autistic-like behaviors did not vary with sex. Data were analyzed by three-way ANOVA with genotype, lighting cycles and sex as variables followed by the Holm-Sidak multiple comparisons’ test. Significant main effects of genotype and lighting, but not of sex, were observed (P< 0.001) (see **Table 3**), however, with a sample size of n=5 per group, per sex, these data sets are not powered to evaluate sex differences, and the negative findings should be viewed with caution. (**B**) Significant main effects of sex were observed on total activity, period, and onset variation (**Table 3**), however, no sex differences were identified between groups. The effects of T7 lighting on circadian rhythms did not differ between males and females of the same genotype. Box plots show the statistical values for each group with the boundary of the box closest to zero indicates the 25th percentile, the line within the box marks the median and the boundary of the box farthest from zero indicates the 75th percentile. The whiskers above and below the box indicate the 90th and 10th percentiles. The sample size was n=6 per group per sex for these measures.

**Fig. 5.**
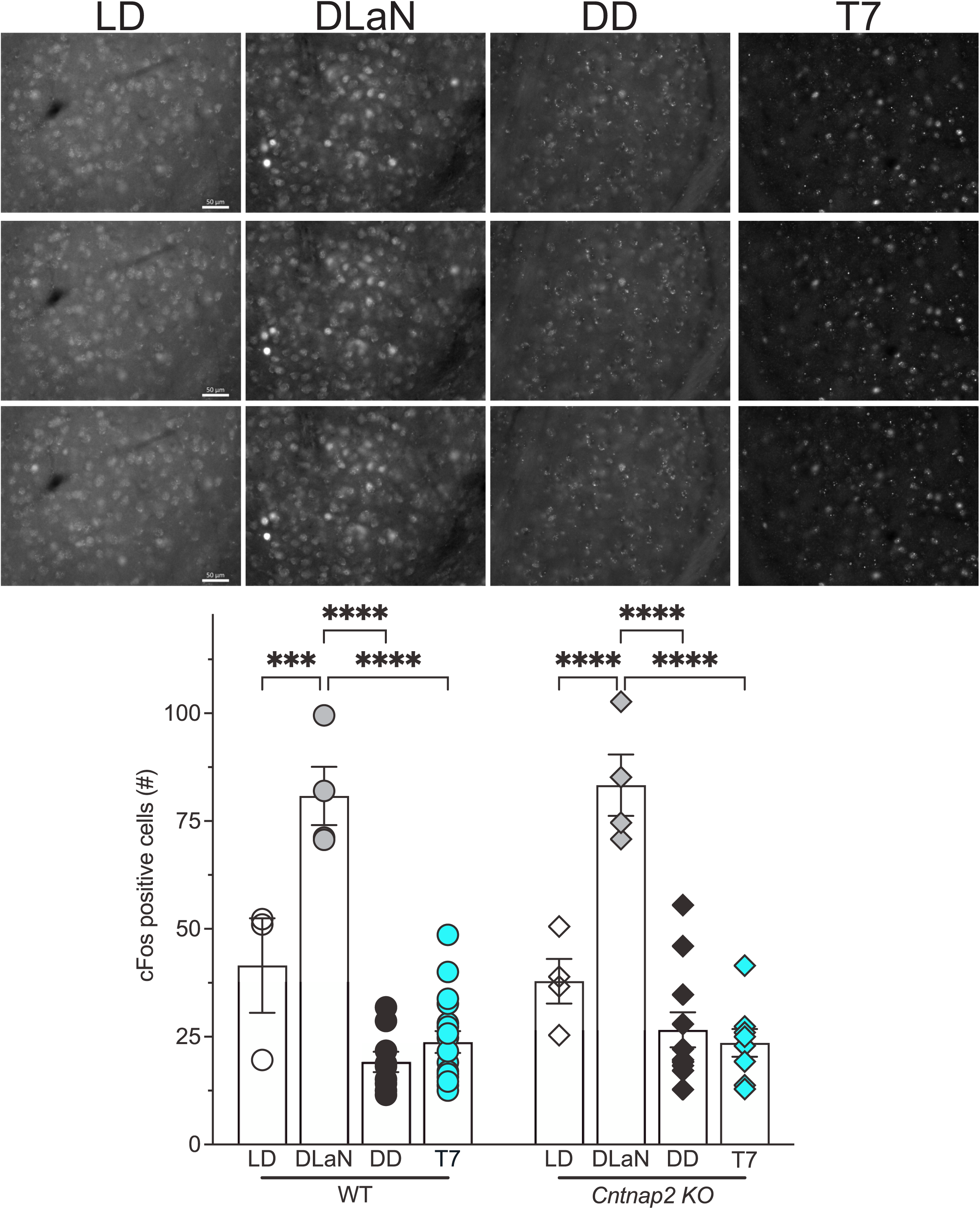
DLaN, but not T7 lighting, evoked an increase in cFos expression in the basolateral amygdala (BLA) of WT and *Cntnap2* KO mice. Mice from each genotype, with a mixed number of males and females, were entrained to a 12h:12h LD cycle for two weeks and then held under one of four lighting conditions: LD, DLaN, DD, and a T7 cycle for two additional weeks (n=3-17 mice per group). Mice were euthanized and perfused at ZT 18 or CT 18, the brains rapidly dissected and processed for immunofluorescence. (**A**) Representative alternate images of cFos expression in the BLA from each of the four groups. Scale bar = 50 µm. (**B**) The number of cFos positive cells was determined bilaterally in the BLA. The values from three to five consecutive coronal sections/animal (both left and right hemisphere) were averaged to obtain one value per animal (n=3 to 17). DLaN elicited a robust increase in the number of cFos expressing cells in both the WT and the mutants; conversely, no induction was observed in any of the other groups. No genotypic differences were observed. Bar graphs show the means ± SD with overlaid the values from the individual WT (circles) and *Cntnap2* KO (diamonds) mice in each of the four lighting conditions: LD (white), DLaN (grey), DD (black), and T7 (teal). Data were analyzed by two-way ANOVA with genotype and treatment as variables (see **Table 4**) followed by the Bonferroni’s multiple comparisons test. The asterisks indicate significant differences lighting conditions within genotype: ***P=0.0002; ****P<0.0001.

**Table 1:**
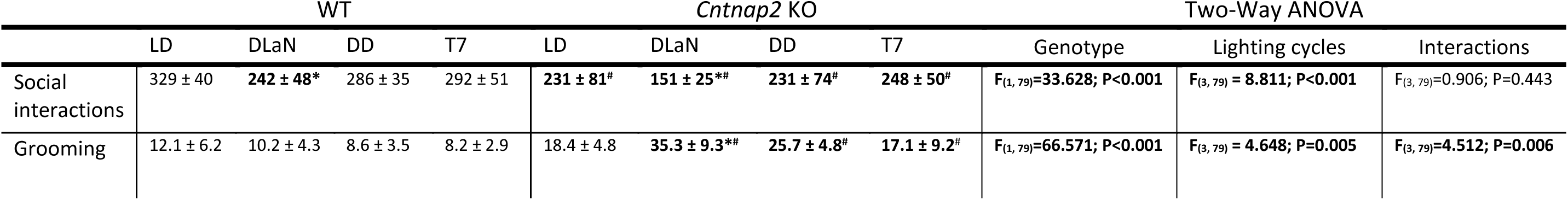
Impact of different lighting cycle conditions on social interactions and grooming in WT and *Cntnap2* KO mice. Values are shown as the averages of the time measured in seconds per each animal per activity ± SD (n= 10 mice per genotype). Data were analyzed by two-way ANOVA with genotype and light cycle conditions as factors, followed by the Holm-Sidäk test for multiple comparisons. Asterisks indicate significant differences between lighting cycles within genotype and crosshatches indicate genotypic differences. Social Interactions: *P=0.006 and 0.002 vs WT and *Cntnap2* KO in LD, respectively; #P<0.05 vs WT in the same lighting cycles. Grooming: *P<0.001 vs *Cntnap2* KO in LD; #P<0.01 vs WT under same lighting cycles. Bolded values indicate significance, alpha= 0.05.

**Table 2:**
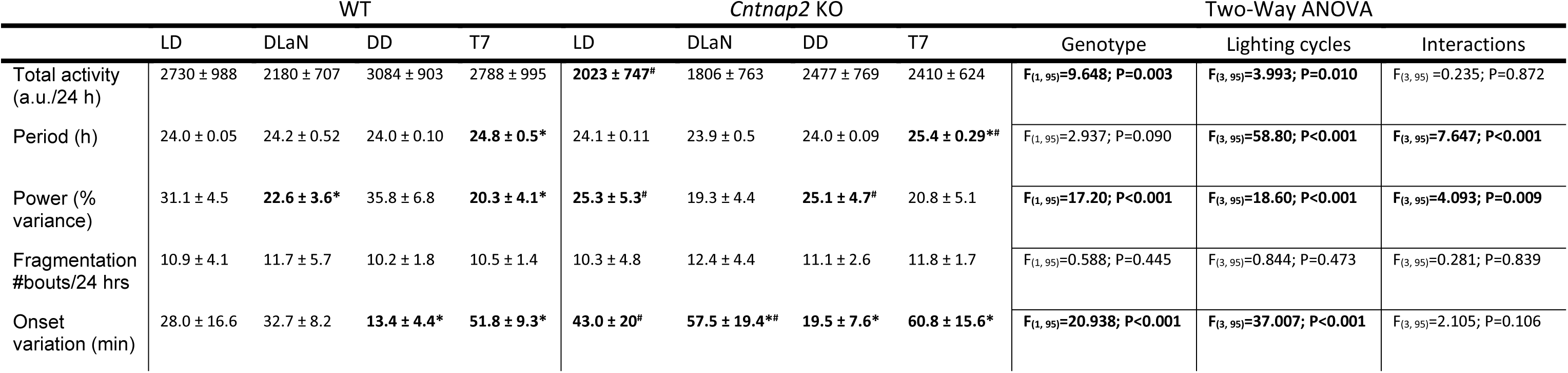
Impact of different light cycle conditions on locomotor activity rhythm parameters in WT and *Cntnap2* KO mice. Values are shown as the averages ± SD (n=12 mice per genotype). Data were analyzed by two-way ANOVA, with genotype and light cycle conditions as factors, followed by the Holm-Sidäk test for multiple comparisons. Asterisks indicate significant differences between light cycles within genotype and crosshatches indicate genotypic differences. *P<0.001 vs WT or *Cntnap2* KO in LD or DD; #P<0.05 vs WT in same light cycles. Bolded values indicate significance, alpha= 0.05.

**Table 3:** Sex affects the impact of different lighting cycle conditions on some parameters of locomotor activity rhythm in WT and *Cntnap2* KO mice. Data were analyzed by three-way ANOVA with sex, genotype, and light cycle conditions as factors, only the values for the interactions among the three values are reported. Values are shown as the averages ± SD (n=6 mice per sex per genotype). Bolded values indicate significance, alpha= 0.05.

|  | Sex | Genotype | Lighting cycles | Interactions |
| --- | --- | --- | --- | --- |
| Total activity (a.u./24 hrs) | **F<sub>(1, 95)</sub>=5.853; P=0.018** | **F<sub>(1, 95)</sub>=10.01; P=0.002** | **F<sub>(3, 95)</sub>=4.144; P=0.009** | F<sub>(3, 95)</sub>=0.648; P=0.586 |
| Period (hrs) | **F<sub>(1, 95)</sub>=13.18; P<0.001** | **F<sub>(1, 95)</sub>=4.239; P=0.043** | **F<sub>(3, 95)</sub>=84.86; P<0.001** | **F<sub>(3, 95)</sub>=2.577; P=0.060** |
| Power (% variance) | F<sub>(1, 95)</sub>=2.441; P=0.122 | **F<sub>(1, 95)</sub>=18.94; P<0.001** | **F<sub>(3, 95)</sub>=20.49; P<0.001** | F<sub>(3, 95)</sub>=0.256; P=0.857 |
| Fragmentation (#bouts/24 hrs) | F<sub>(1, 95)</sub>=1.276; P=0.262 | F<sub>(1, 95)</sub>=0.585; P=0.447 | F<sub>(3, 95)</sub>=0.840; P=0.476 | F<sub>(3, 95)</sub>=0.072; P=0.975 |
| Onset variation (min) | **F<sub>(1, 89)</sub>=8.022; P=0.006** | **F<sub>(1, 89)</sub>=16.508; P<0.001** | **F<sub>(3, 89)</sub>=38.041; P<0.001** | F<sub>(3, 89)</sub>=0.171; P=0.915 |

**Table 4:** DLaN, but not T7, elicits a significant increase in the number of cFos positive cells in the basolateral amygdala (BLA) of WT and *Cntnap2* KO mice. Cells were counted in z-stack images of both the left and right BLA from three to five sections/animal, the values were averaged to obtain one value/section and then one value per animal. Data were analyzed by two-way ANOVA with genotype and lighting cycles as factors followed by the Bonferroni’s multiple comparisons test. Values are shown as the averages ± SD of 3 to 17 mice per genotype and lighting cycle, alpha= 0.05. Bolded values indicate significance. Asterisks indicate significant differences between lighting cycles within genotype; no genotypic differences were observed. *P<0.05 or ***P<0.001 vs WT or *Cntnap2* KO in LD or DD

| Lighting cycle | Genotype |  | Two-Way Anova |  |  |
| --- | --- | --- | --- | --- | --- |
|  | WT | *Cntnap2* KO | Genotype | Lighting cycles | Interactions |
| LD | 41.5 ± 18.9 | 37.9 ± 10.3 | *F*<sub>(1, 52)</sub>=0.1959; P=0.6599 | ***F*<sub>(3, 52)</sub>=60.63; P<0.0001** | *F*<sub>(3, 52)</sub>=0.5808; P=0.6302 |
| DLaN | **80.8 ± 13.5\*\*\*** | **83.3 ± 14.2\*\*\*** |  |  |  |
| DD | **19.1 ± 7.09\*** | 26.6 ± 13.4 | | | |
| T7 | 23.8 ± 10.4 | 23.6 ± 9.05 | | | |

## Results

Difficulty with social interactions is a key symptom of NDD and has been demonstrated in both juvenile (Peñagarikano et al, 2011) and adult (Thomas et al., 2017; Wang et al., 2020; 2023) *Cntnap2* KO mice. Exposure to DLaN exacerbates the social deficits through a mechanism that can be reversed by nightly melatonin treatment or mitigated by shifting to a long-wavelength red light (Wang et al., 2020; 2023). In the present work, social interactions were investigated in WT and *Cntnap2* KO mice held under 4 lighting conditions: LD, DLaN, T7 and DD (**Fig. 1**). While mice under LD and DLaN are synchronized to the environment, their rhythms are not synchronized under T7 and DD conditions. The animals were tested in the three-chamber social arena during their active phase between ZT 16-18 if held in LD or exposed to DLaN, or the subjective night (CT 16-18) for those in DD or T7. The time spent in the chamber with the novel mouse, or the inanimate object was measured (**Fig. 2A**), and significant effects of the factors, genotype and treatment, but not of their interaction, were found by two-way ANOVA (**Table 1**). As previously reported (Wang et al., 2020, 2023), both the WT and mutants were significantly impacted by the DLaN, with the WT exhibiting a reduction in the time spent with the novel mouse under DLaN (P=0.006), while neither T7 (P=0.400) nor DD (P=0.257) altered their social behavior compared to the mice in LD (**Fig. 2A**). Similarly, in the *Cntnap2* KO mice, DLaN treatment reduced the time spent with the novel mouse (P = 0.002) as compared to the mutants in LD, whilst this effect was not observed in mice held in T7 (P=0.879) or DD (P=0.986) (**Fig. 2A**). In comparison to the WT, the *Cntnap2* KO mice exhibited less social interactions in all lighting conditions (LD: P<0.001, DLaN: P<0.001, T7: P=0.049, and DD: P=0.033) (**Fig. 2A**), but only, DLaN, not T7 or DD, reduced social interactions in both genotypes.

Another hallmark symptom of NDDs is repetitive behavior, which is recapitulated in the *Cntnap2* mutants in the form of increased grooming behavior during the day (Peñagarikano et al, 2011; Thomas et al., 2017) and night (Wang et al., 2020; 2023). The mice were tested during the night (ZT 16-18) or subjective night (CT 16-18) and the time spent grooming measured. Significant effects of genotype, treatment, as well as a significant interaction between the two factors were revealed by two-way ANOVA (**Table 1**). In general, the WT mice spent a limited amount of time grooming (**Fig. 2B**), which was not altered by DLaN (P=0.750) or T7 (P=0.339). The *Cntnap2* KO exposed to DLaN exhibited a dramatic increase in grooming (P < 0.001) but a similar augmentation was not seen under T7 (P=0.907) (**Fig. 2B**) as compared to mutants in LD. Compared to WT mice under each condition, the mutants exhibited more grooming under DLaN (P<0.001), T7 (P=0.001) and DD (P<0.01) lighting conditions (**Fig. 2B**), but only DLaN, not T7, exacerbated this aberrant behavior in the mutants.

Prior studies have shown that the *Cntnap2* KO mouse model exhibits disrupted diurnal activity rhythms, including reduced nighttime activity (Thomas et al., 2017; Angelakos et al., 2019; Wang et al., 2020; 2023). To further characterize these disruptions, we assessed locomotor activity rhythms using passive infrared (PIR) sensors under each lighting conditions. At least 10 consecutive days of activity data were collected per animal, and diurnal and circadian parameters were derived. The study was powered to evaluate potential sex differences in activity rhythms (see Wang et al., 2023). As shown by the representative actograms (**Fig. 3A**), both WT and KO mice exhibited rhythms with periods longer than 24 hours when held on the T7 cycles (**Table 2**). Similarly, we observed that both genotypes showed longer free-running periods when assessed by passive infrared recording (PIR) rather than by wheel-running (**Supplemental Fig. 1**; see also Oneda et al., 2022). Significant main effects of genotype and/or lighting condition were identified by two-way ANOVA on total activity, period length, rhythm power and onset variability, along with a significant genotype × lighting interaction for both period and power (**Table 2**).

In WT mice, DLaN significantly reduced the rhythm power (P = 0.001), with no significant changes in other parameters; while the T7 cycles had a broader impact, significantly increasing period (P < 0.001) and onset variability (P < 0.001) whilst also reducing rhythm power (P < 0.001) relative to those in LD and/or DD (**Fig 3B** & **Table 2**). In the *Cntnap2* KO mice, DLaN did not alter the power of the rhythms (P = 0.07) but significantly increased the onset variability (P = 0.037; **Fig 3B**), while T7 illumination significantly lengthened the free-running period (P < 0.001) and increased the variability of the activity onset (P < 0.001; **Table 2** & **Fig 3B**) as compared to their counterpart in LD or DD. Direct comparisons between genotypes revealed significant differences in total activity (P=0.035), rhythm power (P = 0.015) and onset variability (P = 0.004) with no changes in period (P=0.644) under LD, as well as in onset variability (P < 0.001) and period (P < 0.001) under DLaN and T7, respectively. The effect of genotype was less dramatic under DD conditions, where a robust reduction in power (P<0.001) was detected (**Fig**. **3B** & **Table 2**) with no changes in the other parameters. Finally, sex divergent effects were a striking feature of this data set, with significant effects of this variable observed for total activity, period, power and onset variation (three-way ANOVA with genotype, lighting and sex as factors, **Fig.4** & **Table 3**). Overall, the *Cntnap2* KO mice exhibited weaker locomotor rhythms, and these data reveal robust genotype-specific effects of T7 illumination. Furthermore, sex-dependent differences emerged as a prominent feature, underscoring the importance of including both sexes in circadian rhythm studies involving ASD models

Evidence suggests that the BLA, located in the temporal lobe of the cerebral cortex, is a critical mediator of aberrant repetitive behaviors (Felix-Ortiz and Tye, 2014; Sun et al., 2019); in addition, we have previously shown that DLaN elicits increase cFos expression in a population of glutamatergic neurons in this nucleus (Wang et al., 2023). Hence, we examined the expression of this immediate early gene in the BLA of mice held in the 4 lighting conditions (**Fig. 5**). A significant effect of the lighting cycles, but not of genotype, was present (**Table 4**) with both WT and *Cntnap2* KO mice exposed to DLaN displaying a significant increase in the number of cFos positive cells as compared to their counterpart in LD (P=0.0002 and P<0.0001, respectively; **Fig. 5** & **Table 4**); whilst neither the T7 cycle nor DD altered the number of cFos positive cells. In general, both WT and mutants held in T7 or in constant darkness presented with a lower number of cFos positive cells, which was significantly different in WT in DD as compared to those in LD (P=0.0283 WT in DD vs WT in LD). These findings are consistent with the behavioral evidence suggesting that night-time light exposure by itself is not sufficient to activate BLA neurons and elicit the aberrant effects observed.

## Discussion

Patients with NDDs, including ASD, often experience significant disruptions in their sleep-wake cycles. Common complaints include delayed sleep onset, fragmented sleep, and blunted circadian rhythms. We hypothesize that circadian disruption not only contributes to the core symptoms of NDDs but also renders affected individuals more vulnerable to its adverse effects. This vulnerability suggests a potential benefit from circadian-based therapeutic interventions.

To investigate this hypothesis, we employed the *Cntnap2* KO mouse, a well-established model of ASD and related NDDs (Takumi et al., 2020). Our prior work has demonstrated that the *Cntnap2* KO mice exhibit disrupted activity and sleep rhythms, altered neural activity in the SCN, and exaggerated sensitivity to environmental circadian disruption. Notably, exposure to DLaN exacerbates social deficits and repetitive behaviors in these mice (Wang et al., 2020; 2023), the disrupted SCN neural activity and PER2::LUC-driven rhythms in clock gene expression (Wang et al., 2020). Importantly, melatonin administration restores rhythmicity and mitigates behavioral abnormalities, with the greatest improvement observed in animals displaying the most robust circadian rhythms. These findings support a “two-hit” model of NDDs: a genetic predisposition (e.g., *Cntnap2* mutation) primes neural circuits for dysfunction, which can be then amplified by an environmental circadian disruption.

In the current study, we examined the effects of a non-24-hour ultradian lighting schedule— specifically, a T7 cycle (3.5 h light / 3.5 h dark)—on behavior and activity rhythms in *Cntnap2* KO and WT mice. The T7 cycle exposes the animals to light during their active phase each circadian cycle, and both genotypes exhibited free-running rhythms with periods longer than 24 h. Strikingly, *Cntnap2* KO mice did **not** show increased grooming or reduced social interaction under T7, in contrast to their behavioral responses to DLaN (**Fig. 2A,B**), while, consistent with previous studies (Peñagarikano et al, 2011; Thomas et al., 2017; Wang et al., 2020), these behavioral abnormalities persisted under standard LD conditions. Prior work by the Hattar group showed that T7 exposure induces depression-like behaviors (e.g., reduced sucrose preference, increased immobility in the forced swim test) and impairs memory performance in tasks such as the Morris water maze and novel object recognition (LeGates et al., 2012; Fernandez et al., 2018). These impairments were generally assessed during the subjective day. However, recent findings from Fuchs et al. (2023) suggest that behavioral outcomes under T7 conditions are phase-dependent, with mood and memory impairments emerging only when mice are tested during their subjective night.

Although prior reports indicated that the T7 cycle does not overtly disrupt the central circadian clock—based on SCN gene expression, body temperature rhythms, and sleep architecture (Altimus et al., 2008; LeGates et al., 2012)—our high-resolution locomotor activity analysis revealed subtle but important alterations (**Fig. 3**). After two weeks in T7, both WT and KO mice maintained their rhythmicity, albeit with significant lengthening of the period and increased variability in the activity onset, consistent with prior work (Fuchs et al., 2023). Notably, T7 lighting also reduced the power of the rhythms and increased onset variability of the mice relative to DD. Moreover, we detected robust sex differences in these circadian parameters (**Fig. 4**), emphasizing the need to consider sex as a biological variable when evaluating circadian disruption.

Our findingsdings highlight a key dissociation: while T7 alters the activity rhythms, it does not elicit the same behavioral disruptions seen with DLaN in *Cntnap2* KO mice, suggesting that **light at night alone is insufficient** to exacerbate autistic-like behaviors; rather, an interaction between light exposure and circadian misalignment may be required (see also Tam et al., 2021). The differential behavioral outcomes in mice under DLaN vs.T7 provide a valuable framework to dissect mechanistic underpinnings of light-induced pathology.

To identify the relevant neural circuits, we build on previous findings that melanopsin-expressing intrinsically photosensitive retinal ganglion cells (ipRGCs) are necessary for DLaN-induced effects, since *Opn4::DTA* mice lacking these cells do not show DLaN-induced changes in locomotor activity (Wang et al., 2023). Additionally, DLaN increases cFos expression in glutamatergic neurons of the basolateral amygdala (BLA), a region implicated in social behavior and repetitive grooming (Felix-Ortiz and Tye, 2014; Sun et al., 2019). In contrast, T7 exposure did not elicit a cFos response in the BLA (**Fig. 5**), nor did it exacerbate grooming or social deficits. Future studies using chemogenetic silencing or targeted lesions of this BLA cell population will be instrumental in testing its causal role in mediating the behavioral impact of DLaN.

A key innovation of this study is the use of an ultradian T7 lighting schedule to dissociate the direct effects of nighttime light exposure from the circadian disruption it often induces. Prior research has established that DLaN exacerbates behavioral and physiological abnormalities in both WT and *Cntnap2* KO mice. However, it has remained unclear whether these adverse outcomes result from the light exposure per se or from the misalignment of internal circadian rhythms. By implementing a T7 cycle—comprising 3.5-hour light and dark intervals that fall outside the range of circadian entrainment—we provide compelling evidence that the deleterious effects of DLaN are contingent on circadian misalignment. Despite frequent nocturnal light exposure, mutant mice under the T7 cycle maintained their rhythmicity and did not exhibit the worsening of social and repetitive behavior deficits or elevated cFos expression in the amygdala triggered by DLaN. This approach provides a powerful experimental tool to parse light-driven effects from circadian-driven mechanisms and supports the hypothesis that behavioral vulnerability in neurodevelopmental disorder models arises from a two-hit interaction between genetic susceptibility and environmental circadian disruption.

Several limitations should be acknowledged. First, we employed 250 lux illumination for the T7 cycles which was consistent with earlier work. It would be interesting to comparing the behavioral outcomes driven by T7 using different light intensities (250-10 lux) lighting. Second, we measured both the autistic-like behaviors and cFos expression at a fixed phase (ZT/CT 16-18) and did not carry out a time series analysis for either. Of course, given that we have four lighting conditions and two genotypes, a time series sampling every 4 hrs would have required hundreds of additional mice. Third, while our activity rhythm analysis was powered to detect sex differences, this was not the case for the autistic-like behavioral measurements, and the lack of statistically significant sex differences in these measures should be viewed cautiously (**Fig. 4**).

Our findings advance the field by refining the mechanistic understanding of how light at night affects behavior and by introducing ultradian lighting as a novel paradigm for dissecting circadian versus non-circadian influences on neural function. Most importantly, they have translational implications for individuals with NDDs, including ASD, who are frequently exposed to light at nighttime from electronic devices and/or artificial lighting. Our results demonstrate that the adverse behavioral and neural effects of light at night in a genetically susceptible model are contingent on circadian disruption, rather than light exposure alone. This distinction underscores the importance of maintaining circadian rhythmicity as a protective factor against environmental challenges in individuals at risk because of their genetic make-up. Interventions aimed at supporting circadian health—such as structured light exposure schedules, consistent sleep-wake timing, or the use of circadian-stabilizing agents like melatonin—may help mitigate the impact of nocturnal light exposure in individuals with NDDs/ASD. Furthermore, the ultradian lighting paradigm used here provides a mechanistically informative model for testing the efficacy of such circadian-targeted interventions without introducing the confounding effects of arrhythmicity. Ultimately, these findings support the development of chronobiology-informed strategies to improve behavioral outcomes and quality of life in individuals with NDDs.

## Acknowledgments

We would like to thank Derek Dell’Angelica for his help with the BioRender software and Alina Ryu who helped with the scoring of the fluorescence staining. Dr. Kathy Tamai maintained the mouse colonies used for the generation of the mice and participated in discussions of the work.

## Conflicts of Interest

The authors declare no conflicts of interest.

## Author contributions

SAMV, HBW, GDB, CAG and CSC conceived the hypothesis and experimental design of this study. SAMV, HBW, performed the behavioral experiments and analysis. SAMV, KNN, CTC, GS carried out the immunofluorescence and analysis. CTC assisted with literature review and referencing. SAMV and HBW wrote the first draft; GDB, CAG and CSC edited, wrote and compiled the final version paper with contribution from the other authors.

## Data Sharing

Available from the corresponding author upon request.

---

**Supplemental Fig. 1:** The illuminance of the LED lighting was 250 lx as measured from the floor of the animal holding chamber. The irradiance was 75 microW/cm2 with a peak at 450nm and a Melanopic to Photopic (M/P) ratio of 0.57. The M/P ratios were calculated using the rodent circadian lighting toolbox courtesy of Dr. S. Peirson (Sleep and Circadian Neuroscience Institute, Oxford, UK).

**Supplemental Fig. 2:** Free-running period was measured in WT and *Cntnap2* KO mice under constant darkness, with either running wheels or passive infrared (PIR) sensors. Each dot represents an individual animal. WT mice exhibited a significantly longer circadian period when assessed by PIR compared to wheel-running. A similar effect of measurement method was observed in KO mice. The same recording chambers were used to measure activity with both wheels and PIRs. The activity rhythms shown were measured from two distinct cohorts of WT mice in 2019 and 2020, respectively. The data from the 2020 cohorts were analyzed using two-way ANOVA, which found significant effects of the type of apparatus used to record the rhythms in activity (F = 28.393, P < 0.001) but not genotype (F = 1.361, P = 0.253). These findings highlight the influence of the method used for measurement of circadian period.

## Notes

Funding: Supported by Eunice Kennedy Shriver National Institute of Child Health and Human Development under award number: 5U54HD087101 (PI S. Bookheimer, H. Kornblum). GDB received University of California funding.

### Competing Interest Statement

The authors have declared no competing interest.

