## Supplemental figures for "Dissociating the Effects of Light at Night from Circadian Misalignment in a Neurodevelopmental Disorder Mouse Model Using Ultradian Light-Dark Cycles"

### Supplemental Fig. 1

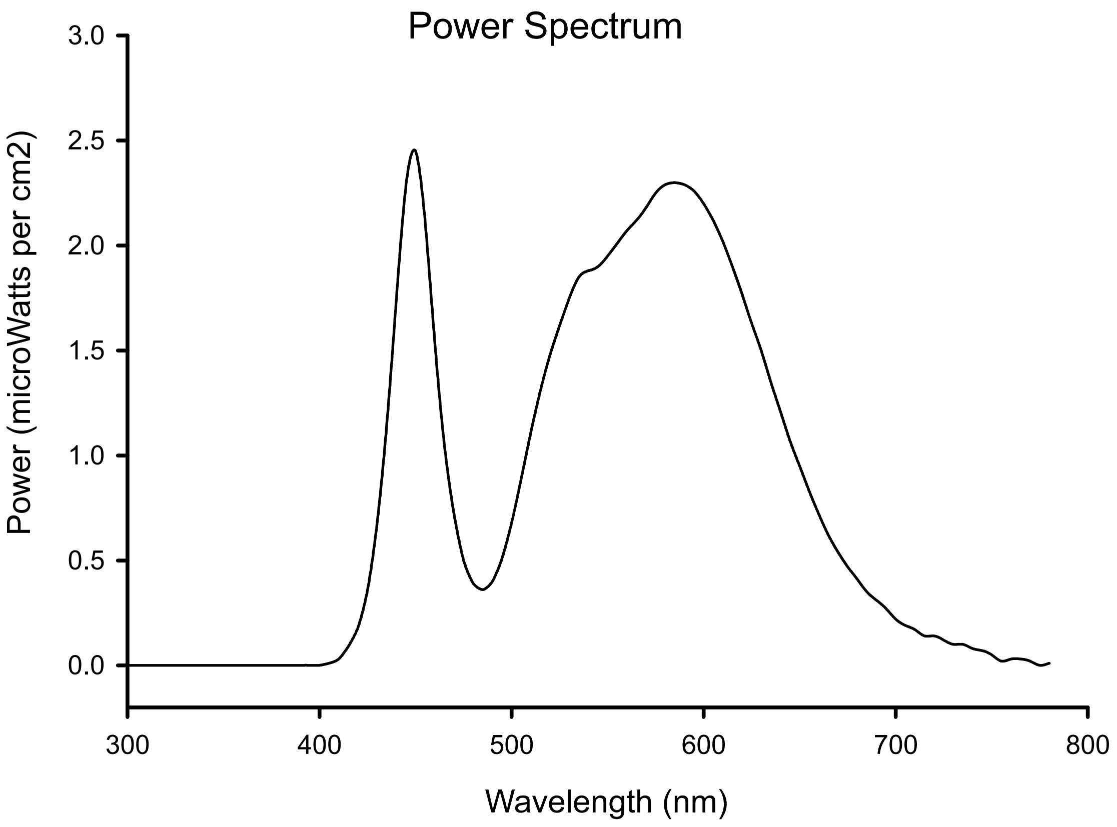


Supplemental Fig. 1: The illuminance of the LED lighting was 250 lx as measured from the floor of the animal holding chamber. The irradiance was 75 microW/cm2 with a peak at 450nm and a Melanopic to Photopic (M/P) ratio of 0.57. The M/P ratios were calculated using the rodent circadian lighting toolbox courtesy of Dr. S. Peirson (Sleep and Circadian Neuroscience Institute, Oxford, UK).

### Supplemental Fig. 2

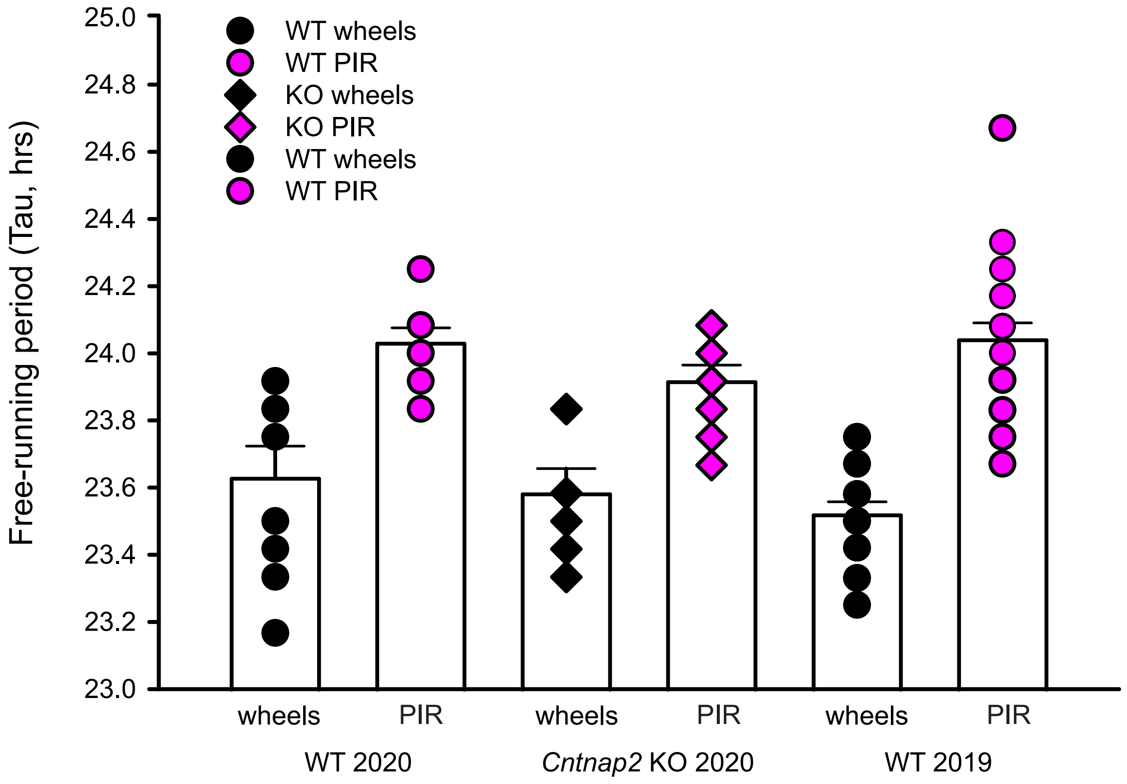


Supplemental Fig. 2: Free-running period was measured in WT and Cntnap2 KO mice under constant darkness, with either running wheels or passive infrared (PIR) sensors. Each dot represents an individual animal. WT mice exhibited a significantly longer circadian period when assessed by PIR compared to wheel-running. A similar effect of measurement method was observed in KO mice. The same recording chambers were used to measure activity with both wheels and PIRs. The activity rhythms shown were measured from two distinct cohorts of WT mice in 2019 and 2020, respectively. The data from the 2020 cohorts were analyzed using two-way ANOVA, which found significant effects of the type of apparatus used to record the rhythms in activity ($F = 28.393$, $P < 0.001$) but not genotype ($F = 1.361$, $P = 0.253$). These findings highlight the influence of the method used for measurement of circadian period.
